# MMP21 behaves as a fluid flow transported morphogen to impart laterality during development

**DOI:** 10.1101/2024.10.13.618030

**Authors:** Tim Ott, Amelie Brugger, Emmanuelle Szenker-Ravi, Yvonne Kurrle, Olivia Aberle, Matthias Tisler, Martin Blum, Sandra Whalen, Patrice Bouvagnet, Bruno Reversade, Axel Schweickert

## Abstract

Heterotaxy (HTX) is frequently caused by deleterious variants in the gene encoding Matrix metallopeptidase 21 (MMP21). However, the underlying pathomechanism has not been ascertained. In this study, we report on a novel HTX-associated *MMP21* knockout allele in humans and investigate the peptidase’s role during laterality development using *Xenopus* embryos as animal model. The targeted inactivation of *mmp21* in f0 mutant *Xenopus* successfully phenocopied the human HTX condition, yet the cilia-driven leftward fluid flow, which initiates asymmetric gene activity at the left-right organizer (LRO), was unaltered in *mmp21* null frogs. Instead, our analysis of downstream events revealed that flow response, the left-sided repression of *dand5*, could not take place. Remarkably, gain-of-function experiments demonstrated that Mmp21 spreads over LRO cells and triggers flow response. Additionally, Mmp21 functions upstream of Cirop, another metallopeptidase, which we found specifically localized to LRO cilia. Thus, our findings suggest that Mmp21 may be the long-sought morphogen, which is actively transported by the leftward fluid flow to Cirop-laden cilia, in order to specify the left side of the embryo.

## Introduction

The molecular definition of the left-right body axis is a crucial step during early embryogenesis, allowing complex chiral organs like the vertebrate heart to orderly emerge and to be asymmetrically positioned within body cavities. In humans, the normal arrangement of thoracic and abdominal organs is called *situs solitus* (SS), whereas *situs inversus* (SI) describes a complete mirror-imaged orientation, not necessarily associated with health constraints. In contrast, severe health conditions can occur if single or multiple organs are misoriented, multiplied or missing, commonly referred to as heterotaxy (HTX)^1^. Estimations for the frequency of laterality defects are difficult to make as methodologies and definitions vary between epidemiological studies in the field. However, a retrospective 12-year spanning population-based study of infants diagnosed with syndromic or non-syndromic laterality defects determined an overall prevalence of 1.9 per 10 000 live births^2^.

Humans and all other vertebrates, except Sauropsida and Cetartiodactyla, utilize a cilia-dependent mechanism to establish laterality^3,4^. During neurulation, a ciliated epithelium, called left-right organizer (LRO) arises at the embryonic midline^4^. Central LRO cells harbor posteriorly polarized motile mono-cilia, which rotate in a clockwise fashion thereby generating a transient leftward flow of extracellular fluid^5,6^. The fluid flow represents the initial asymmetric stimulus, which is exclusively perceived on the left side by lateral LRO cells through sensory cilia^7,8^. These sensor cells express a specific set of morphogens, most importantly and archetypically: Nodal homolog 1 (Nodal1) and its antagonist DAN domain BMP antagonist family member 5 (Dand5)^9^. Flow response post-transcriptionally downregulates left-sided *dand5* expression^10–12^. Hence, Nodal1 is released from repression^13^ and travels from the midline to the left lateral plate mesoderm (LPM) to activate the so-called Nodal signaling cascade^14^. This cascade is conserved among all vertebrates and involves self-induction of a Nodal homolog, allowing its propagation throughout the left LPM^15^. At the same time, Nodal signaling induces the feedback inhibitor Left-right determination factor (Lefty) as well as the transcription factor Paired like homeodomain 2 (Pitx2). Whereas Lefty restricts the spatio-temporal action of Nodal ligands^16^, Pitx2 eventually directs asymmetric organogenesis^17^.

In 2015, Matrix metallopeptidase 21 (MMP21) was discovered as a novel laterality determinant in patients with deleterious *MMP21* variants, causing a form of autosomal recessive HTX (MIM 616749)^18–20^. Loss of Mmp21 in mice^18,20^ and zebrafish^19,20^ resembled the human left-right patterning defects, indicating a highly conserved role in the lateralization process. Although it was reported that cilia motility in the murine LROs was not disturbed in *Mmp21* mutants^18^, the hierarchical placement and function of Mmp21 remained otherwise elusive. Remarkably, we recently found that *MMP21* belongs to an ancient genetic module, which is only preserved in vertebrates with a fluid flow generating LRO^3^.

Here we report on a novel HTX-associated germline variant of *MMP21* and dissect the role of Mmp21 during symmetry breakage leveraging on the power of the *Xenopus* model system^21^. Through loss-of-function (LOF) experiments, we show that Mmp21 is situated in a hub downstream of flow but upstream of left-sided *dand5* repression. Gain-of-function (GOF) experiments reveal that Mmp21 acts as a non-cell-autonomous fluid flow effector, which is able to travel over LRO cells following its secretion. Moreover, our study demonstrates that Mmp21 operates upstream of another peptidase called Ciliated left-right organizer metallopeptidase (Cirop). *CIROP*, like *MMP21*, is associated with HTX (MIM 619702) and belongs to the same fluid flow-related gene module^3^. As Cirop localizes to cilia of LRO sensor cells, we suggest that Mmp21 is a morphogen, which diffuses with the leftward fluid flow to specify the left side in a Cirop-dependent manner.

## Results

### The domain architecture of human MMP21 is conserved in *Xenopus*

MMP21 and all other MMP family members are zinc-dependent endopeptidases of the metzincin type (Fig. 1A). This type is characterized by the presence of a HEXXH motif in the catalytic domain (CD) followed by a methionine-turn (Met-turn) in close proximity. The glutamate of the HEXXH motif exerts catalytic function, whereas the two histidines together with a more C-terminal histidine, glutamic or aspartic acid serve as zinc ligands. MMPs show an extended consensus motif (HEXXHXXGXXH) and are synthesized as preproproteins. In latent MMPs, a conserved cysteine within the propeptide binds the catalytic zinc atom to block enzymatic activity, a mechanism known as cysteine switch. MMP21 has a Furin cleavage site (FCS) as well as C-terminal Hemopexin-like repeats, which create a four-bladed beta-propeller, a structure known to provide substrate specificity and to mediate inhibitor interaction in other MMPs^22^. These features are conserved in *Xenopus* Mmp21 (Fig. 1B) and fold into a protein that, after maturation (Fig. S1A), is predicted by ColabFold^23^ to possess a bipolar, globular structure with an open catalytic cleft (Fig. S1B).

**Fig. 1:**
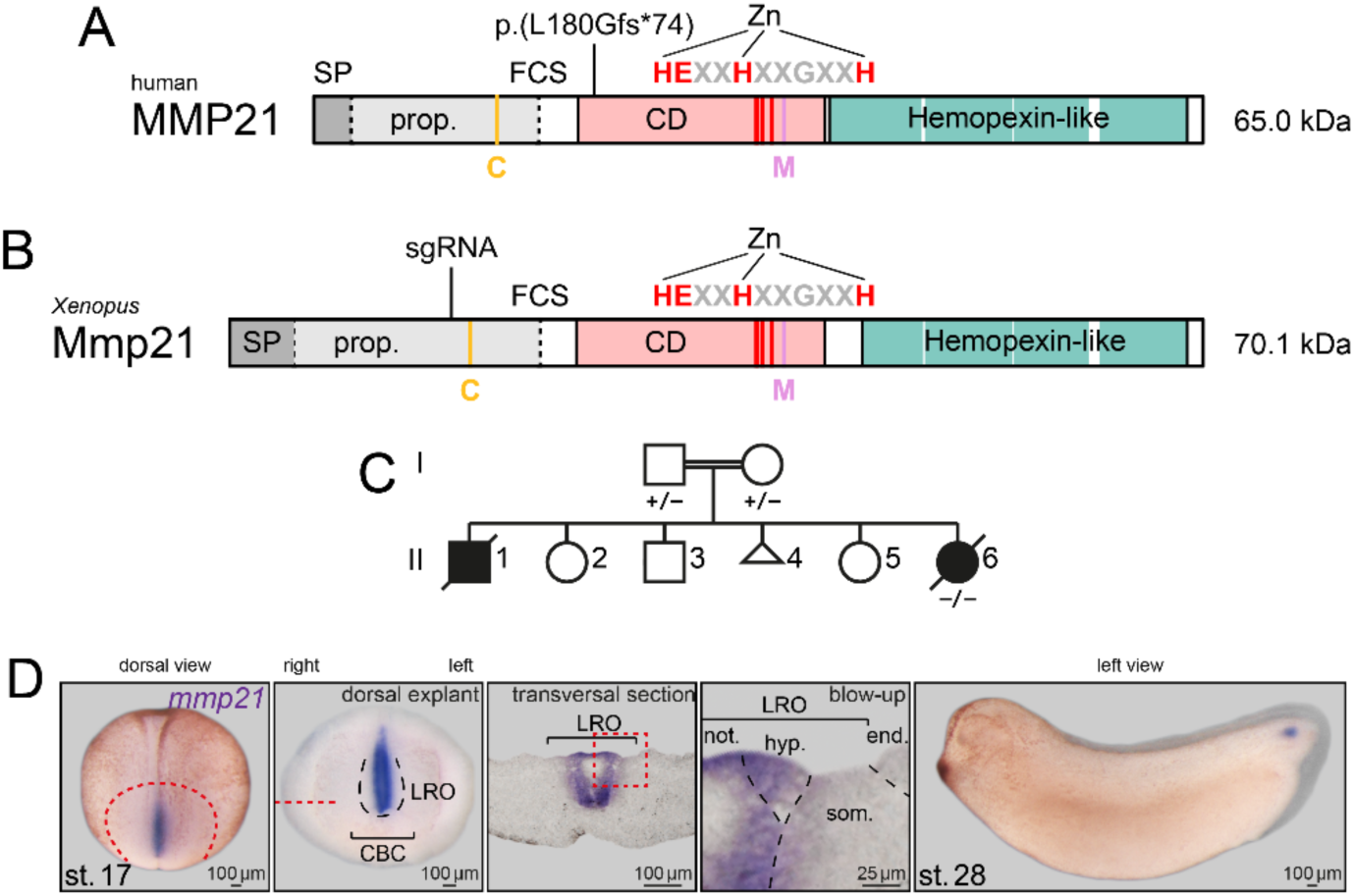
The homozygous variant p.(L180Gfs*74) of LRO-associated *MMP21* causes laterality defects. **(A)** MMP21 consists of an N-terminal signal peptide (SP), a propeptide with Furin cleavage site (FCS), a catalytic domain (CD) and C-terminal Hemopexin-like repeats. Zn coordination as well as conserved residues of the cysteine-switch (C), catalytic core (HEXXHXXGXXH) and methionine-turn (M) are marked. The position of the frameshift mutation p.(L180Gfs*74) is indicated in the CD. **(B)** Domain composition of *Xenopus* Mmp21 is equivalent to that of human MMP21. **(C)** The pedigree of the female proband (II-6) illustrates that both she and the youngest sibling (II-1) presented with heterotaxy (HTX). Wild-type and mutant *MMP21* alleles are indicated by a plus and minus symbol, respectively. **(D)** During *Xenopus* development, *mmp21* is expressed in the central flow-generating cells of the LRO with noto- and hypochordal fate. The sensory LRO cells with somitic fate, surrounded by endodermal cells, as well as the circumblastoporal collar (CBC), are free of *mmp21* transcripts. In later stages, expression of *mmp21* is restricted to the tailbud and successively vanishes. Dashed red lines indicate consecutive incision and transversal sectioning. Dashed red line square outlines the magnified region. Dashed black lines display tissue borders.

### A novel biallelic frameshift variant of *MMP21* causes HTX

The relevance of MMP21 for normal development is already evident but was further emphasized by our finding of a novel *MMP21* null allele in a cohort of 108 index cases with laterality defects (ID 4263)^3^. Exome sequencing revealed a biallelic germline deletion c.537_547delGCTGCTGGGCG in the female proband (II-6) with familial HTX, born to consanguineous parents (Fig. 1C). This small deletion translates to an early frameshift variant p.(L180Gfs*74), which is suspected to induce nonsense-mediated mRNA decay (Fig. 1A).

### *Xenopus mmp21* is specifically expressed in central LRO cells

To get an entry point for our functional study, we then analyzed the spatio-temporal activity of *mmp21* via RNA ISH in *Xenopus laevis* (Fig. 1D). *mmp21* expression starts during neurulation and is pronounced when leftward fluid flow peaks at stage 17. Transcripts appear in posterior cells of the embryonic midline with hypo- and notochordal fate. These cells include the central flow-generating cells of the LRO but not the somitic LRO sensor cell population. After neurulation, the expression of *mmp21* gradually diminishes, though it remains detectable in the tailbud during later stages. Notably, *mmp21* expression in zebrafish embryos is restricted to LRO cells as well^20^, suggesting a specific role for the peptidase in events occurring within that tissue.

### Disturbed flow response induces laterality defects in *mmp21* mutants

In order to characterize the physiological function of Mmp21, we chose a CRISPR/Cas9 based LOF approach to generate f0 mutants, also known as crispants. Zygotes were injected with Cas9 ribonucleoprotein (CRNP) made with a sgRNA directed against the second exon of *mmp21* (Fig. S2A). This target site corresponds to D142 within the propeptide of Mmp21 (Fig. 1B). Direct sequencing of PCR products^24^ from pooled genomic DNA of stage 45 tadpoles and subsequent Synthego Inference of CRISPR Edits (ICE) analysis confirmed efficient mutagenesis of the *mmp21* locus with an indel rate of 92 % (Fig. S2A). *mmp21* crispants presented high rates of HTX and SI, when scored for heart, gallbladder and intestine orientation at stage 45, but developed otherwise normally (Fig. 2A), thereby phenocopying humans with biallelic *MMP21* LOF variants.

**Fig. 2:**
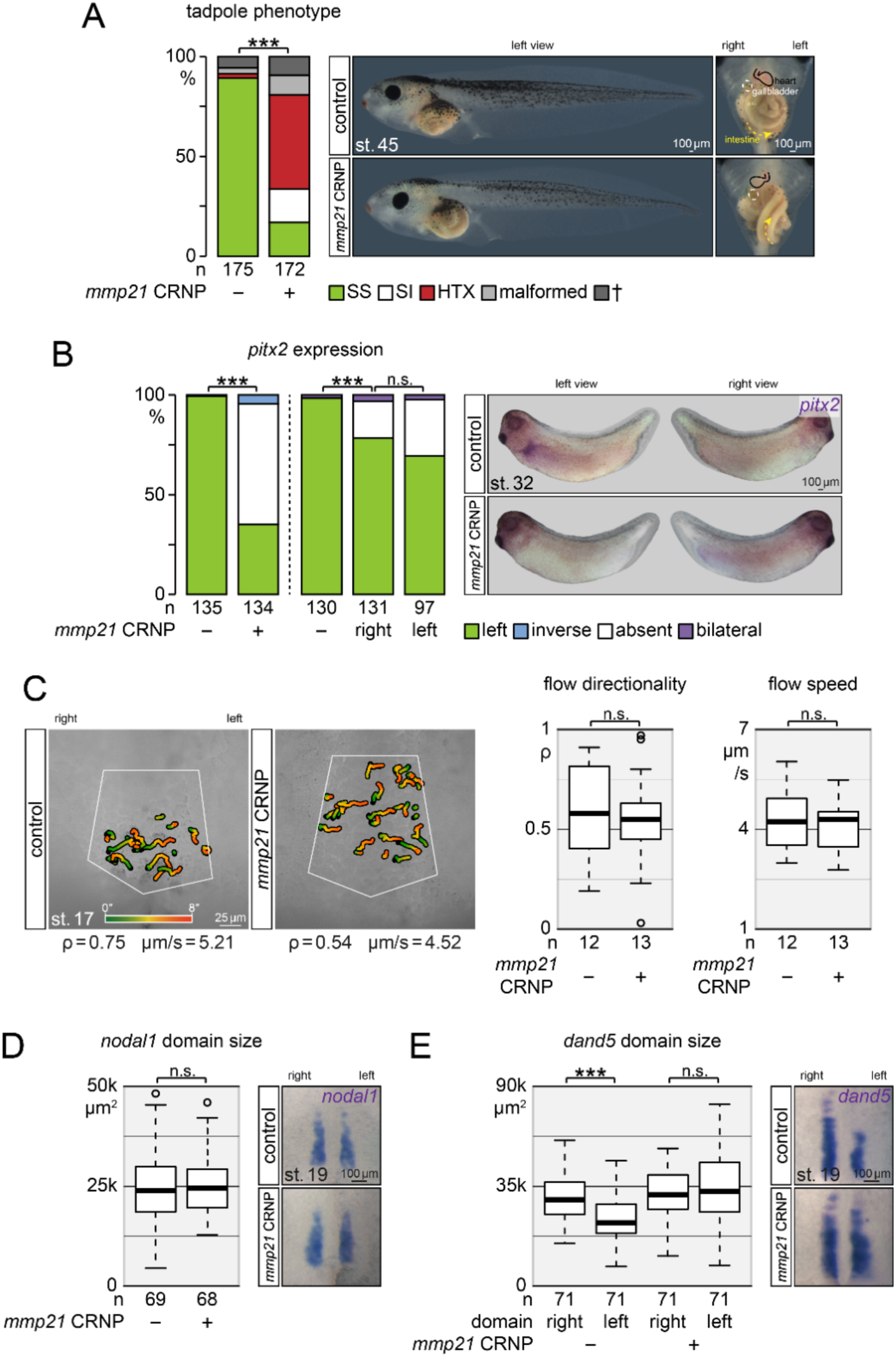
Loss of Mmp21 disturbs laterality owing to disturbed fluid flow response. **(A)** Genome editing of *mmp21* through Cas9 ribonucleoprotein (CRNP) injections interferes with *situs solitus* (SS) and induces high rates of *situs inversus* (SI) and heterotaxy (HTX). Crispant embryos developed otherwise normally with low mortality and malformation rates. **(B)** Left-sided *pitx2* expression is absent in Mmp21 depleted embryos. Uniquely, sided-injections revealed a bilateral requirement of Mmp21. **(C)** Speed and directionality of the ciliary driven leftward fluid flow was not affected by a loss of Mmp21. Color-graded gradient time trails depict the displacement of fluorescent microspheres, which were used for analysis, within an 8 s time window in the white-outlined LRO region. **(D)** The *nodal1* domain at the LRO was not reduced in size in *mmp21* mutants, but **(E)** flow-dependent downregulation of *dand5* on the left side was compromised.

Assessments of the Nodal cascade in Mmp21 depleted embryos were conducted with a *pitx2* ISH, which revealed absence of left-sided cascade activity (Fig. 2B). Of note, reintroduction of *mmp21* mRNA rescued *pitx2* expression in the left LPM, thus demonstrating specificity of the LOF phenotype (Fig. S2B). Surprisingly, when we mutagenized *mmp21* exclusively on the left or on the right side, we found both sides contributing to the phenotype (Fig. 2B), a characteristic that was never reported for a factor in this context before.

Due to our observation that *mmp21* is expressed in the central proportion of the LRO, a cell population only known for the generation of the leftward fluid flow, we analyzed fluid flow dynamics of *mmp21* mutants via videomicroscopy-based tracking of applied fluorescent microspheres (Fig. 2C). However, neither flow directionality nor flow speed was disturbed, in good agreement with the undisturbed cilia motility in *Mmp21* mutant mice^18^.

In the light of these results, we hypothesized that loss of Mmp21 should manifest at the level of LRO sensor cells, which perceive and interpret the leftward fluid flow. Hence, LRO *nodal1* and *dand5* expression domains were quantified post-flow in *mmp21* crispants using ISH and subsequent image thresholding (Fig. 2D & E). As expected, the domain size of *nodal1* was unaltered in *mmp21* mutants. Intriguingly, comparison of left versus right *dand5* domains unveiled that regular left-sided attenuation of *dand5* mRNA level, i.e. flow response, failed in Mmp21-depleted embryos. Accordingly, *pitx2* expression was reestablished in *mmp21* crispants by a left-sided translation-blocking morpholino oligomer (TBMO) mediated knockdown of *dand5* to mimic flow response (Fig. S2C). These results suggest that downstream events like Nodal1 maturation, secretion, transfer to the LPM or self-induction are Mmp21-independent processes. Thus, Mmp21 function is situated in a hub downstream of flow but upstream of *dand5* repression.

### Mmp21 triggers the Nodal-cascade non-cell-autonomously

Our initial rescue experiments showed a relatively high proportion of embryos that activated the Nodal cascade not only on the left but also ectopically on the right side, raising the question whether this phenotype is representative for a Mmp21 GOF. To test this idea, we overexpressed Mmp21 in left versus right LRO cells followed by a *pitx2* ISH (Fig 3A). Mmp21 was found to be a potent inducer of bilateral Nodal signaling, particularly when targeted to the right side. However, left-sided *mmp21* injections were also able to initiate the cascade on the contralateral side but with reduced efficacy. Strikingly, frequencies of virtually 100 % could be achieved with the same left-sided injection scheme by gradually increasing the *mmp21* dose, indicating that Mmp21 acts non-cell-autonomously. Ectopic activation of the Nodal cascade can result from a defective midline barrier, which is a transient group of axial cells expressing Lefty to shield the right LPM from left-sided Nodal-signaling^25^. To exclude that Mmp21 affects this structure, we first analyzed *lefty* expression along the midline with ISH in embryos overexpressing Mmp21, but found no reduction on mRNA level (Fig. S3A). Since Mmp21 and Lefty may interact on protein level and because one would expect a delayed right-sided cascade initiation if this phenotype was caused from midline leakage, we monitored the onset of *nodal1* expression stage-wise with an ISH (Fig. S3B). Within this fine-meshed temporal resolution, we observed a synchronized emergence of left- and right-sided *nodal1* expression in Mmp21 GOF embryos. Therefore, we conclude that Mmp21 is a non-cell-autonomous inducer of the Nodal cascade, which does not compromise the barrier function of the midline.

**Fig. 3:**
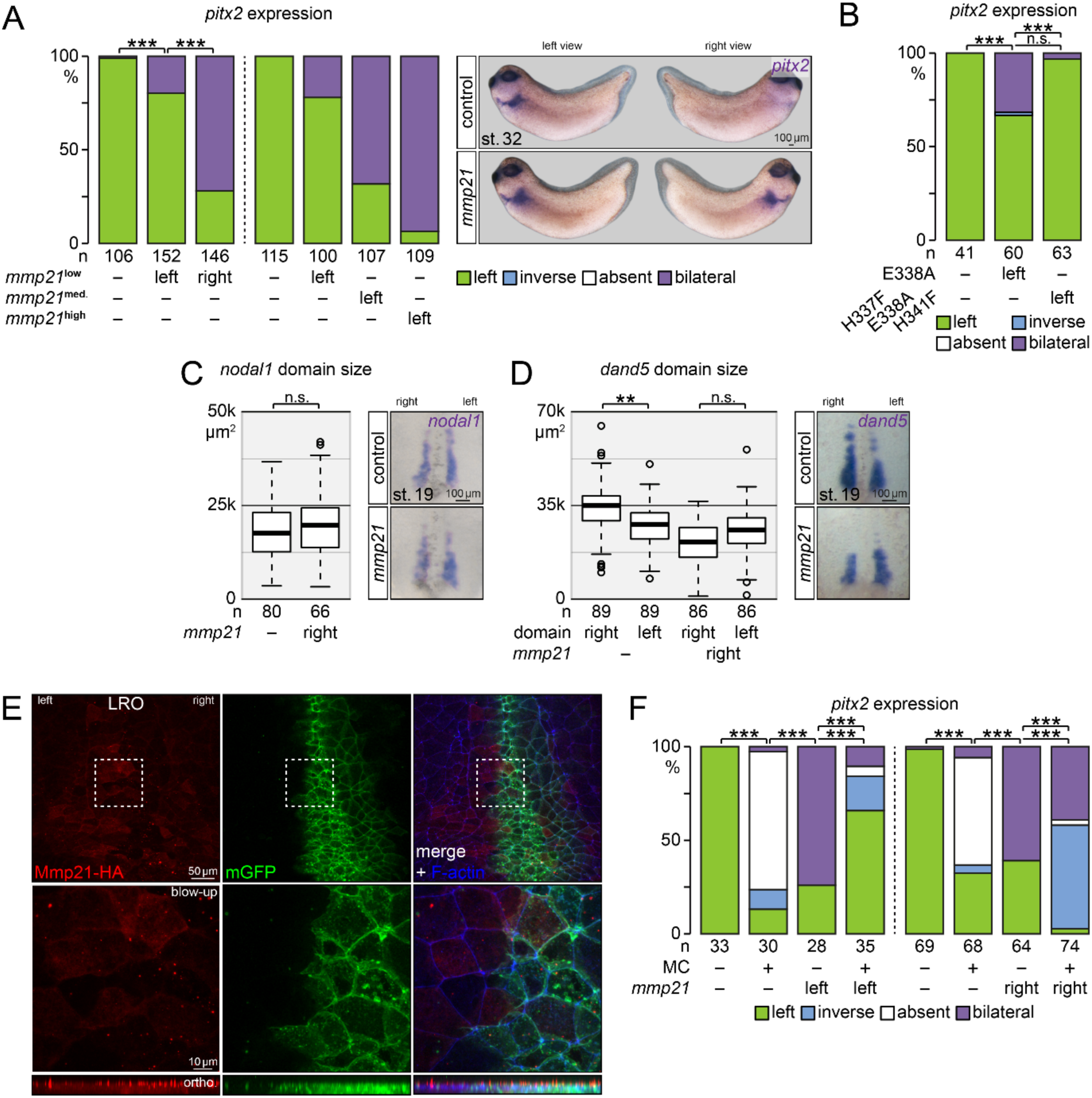
Mmp21 behaves as a morphogen and acts as fluid flow effector. **(A)** Unilateral injections of *mmp21* mRNA induce ectopic expression of *pitx2* on the right side. This effect was more pronounced on the right side, although higher *mmp21* mRNA doses compensate for the difference. **(B)** The active site mutant (E338A) but not the full HEXXH motif mutant (H337F-E33A-H341F) Mmp21 retained the Nodal-cascade inducing effect. **(C)** *nodal1* expression was not impacted by overexpressed Mmp21, whereas **(D)** the right *dand5* domain was reduced in size to the level of the left side post-flow. **(E)** C-terminally HA tagged Mmp21 spreads over LRO cells. Co-expressed mGFP marks targeted cells on the right side. F-actin staining indicates cell borders. Dashed white line squares outline the magnified regions, highlighting Mmp21 presence on targeted (expressing) and non-targeted (non-expressing) cells. Orthogonal view shows enrichment of Mmp21 above apical mGFP and F-actin signals. **(F)** Methylcellulose (MC) increases the viscosity of the extracellular fluid and restricts the field of action of overexpressed and endogenous Mmp21, as judged by the expression pattern of *pitx2*.

Next, we wanted to assess if the catalytic activity of Mmp21 is necessary for its function (Fig. 3B). Surprisingly, an active site mutant E338A construct was still able to induce the Mmp21 GOF phenotype. In contrast, the triple mutant H337F-E338A-H341F construct, in which the two HEXXH domain histidines were additionally changed to phenylalanine, lost the potency to induce the Nodal cascade. These data imply that the integrity of the CD is likely required for Mmp21 function, whereas its enzymatic activity might not be crucial^26,27^.

Our LOF analysis showed that Mmp21 is required for the downregulation of *dand5* in left LRO sensor cells. Consequently, we hypothesized that the gain of Mmp21 attenuates *dand5* in right sensor cells, too, thereby inducing ectopic Nodal signaling. In agreement with this idea, *nodal1* expression at the LRO was not affected by overexpressed Mmp21 (Fig. 3C) but *dand5* on the right side was reduced to the level of the left side post-flow (Fig. 3D).

### Mmp21 acts as a morphogen

Expression of *mmp21* in central LRO cells, Mmp21-dependent *dand5* repression in LRO sensor cells, non-cell-autonomous induction of the Nodal cascade by Mmp21 and the presence of a N-terminal signal-peptide in Mmp21 pointed to a scenario in which the peptidase specifies the left-right axis as would a classical morphogen. Hence, we decided to analyze the localization of Mmp21 via immunofluorescence (Fig. 3E). Visualization was achieved with an HA-tagged Mmp21, as one commercial and two custom-made antibodies failed to detect the endogenous protein. Mmp21-HA, which was unilaterally expressed in LRO cells, showed a diffuse distribution with scattered apical aggregates. Remarkably, Mmp21-HA was not limited to targeted cells marked by co-injected mGFP but spread over the LRO as would be expected for a diffusible signaling molecule. To verify that the observed distribution is not artifactual, injected embryos were incubated during LRO formation with Brefeldin A, a fungal lactone known to block vesicle transport along the endoplasmic reticulum to Golgi route, thus inhibiting secretion^28^. Brefeldin A effectively reduced the spreading of Mmp21, limiting it to the injected mGFP-positive LRO cells (Fig. S3C). Notably, the diffusion of Mmp21 was also observed if the construct was targeted to LRO-unrelated naïve epithelial cells of the animal cap (Fig. S3D). In this tissue, Mmp21 was not only found in vesicular structures of targeted, mGFP-positive cells but also in clusters along the apical cell borders of non-targeted, mGFP-negative cells, which resembled the localization of Wnt and Hedgehog ligands during their diffusion^29,30^.

In order to study if extracellular mobility is relevant for Mmp21 function, we utilized archenteron injections of methylcellulose (MC), a minimally invasive method to increase the viscosity of the extracellular fluid over the LRO (Fig. 3F)^31^. The MC injections alone prevented the induction of the Nodal cascade. Notably, MC also prevented the non-cell-autonomous contralateral induction of the Nodal cascade caused by left-sided *mmp21* injections, which was sufficient to restore wild-type Nodal signaling in an autocrine fashion. Likewise, the Nodal cascade was inverted if MC was injected into embryos that overexpressed Mmp21 on the right side. In conclusion, MC restricts the field of Mmp21 action to expressing cells, reinforcing the notion that the peptidase may acts as a *bona fide* morphogen in the context of leftward fluid flow.

### Cirop is a downstream effector of Mmp21 on LRO cilia

We recently identified and characterized Cirop, another peptidase of the metzincin type (Fig. 4A), which is also required downstream of flow but upstream of *dand5* repression. However, unlike Mmp21, Cirop harbors a C-terminal transmembrane domain, is expressed in LRO sensor cells, acts cell-autonomously on the left side and is unable to induce HTX upon overexpression^3^. These findings suggested to us that Cirop constitutes a permissive signal for left-right axis specification and could therefore function as a downstream effector of Mmp21. This conception was further supported by the striking cilia localization of HA-tagged Cirop in targeted mGFP-positive LRO sensor cells (Fig. 4B).

**Fig. 4:**
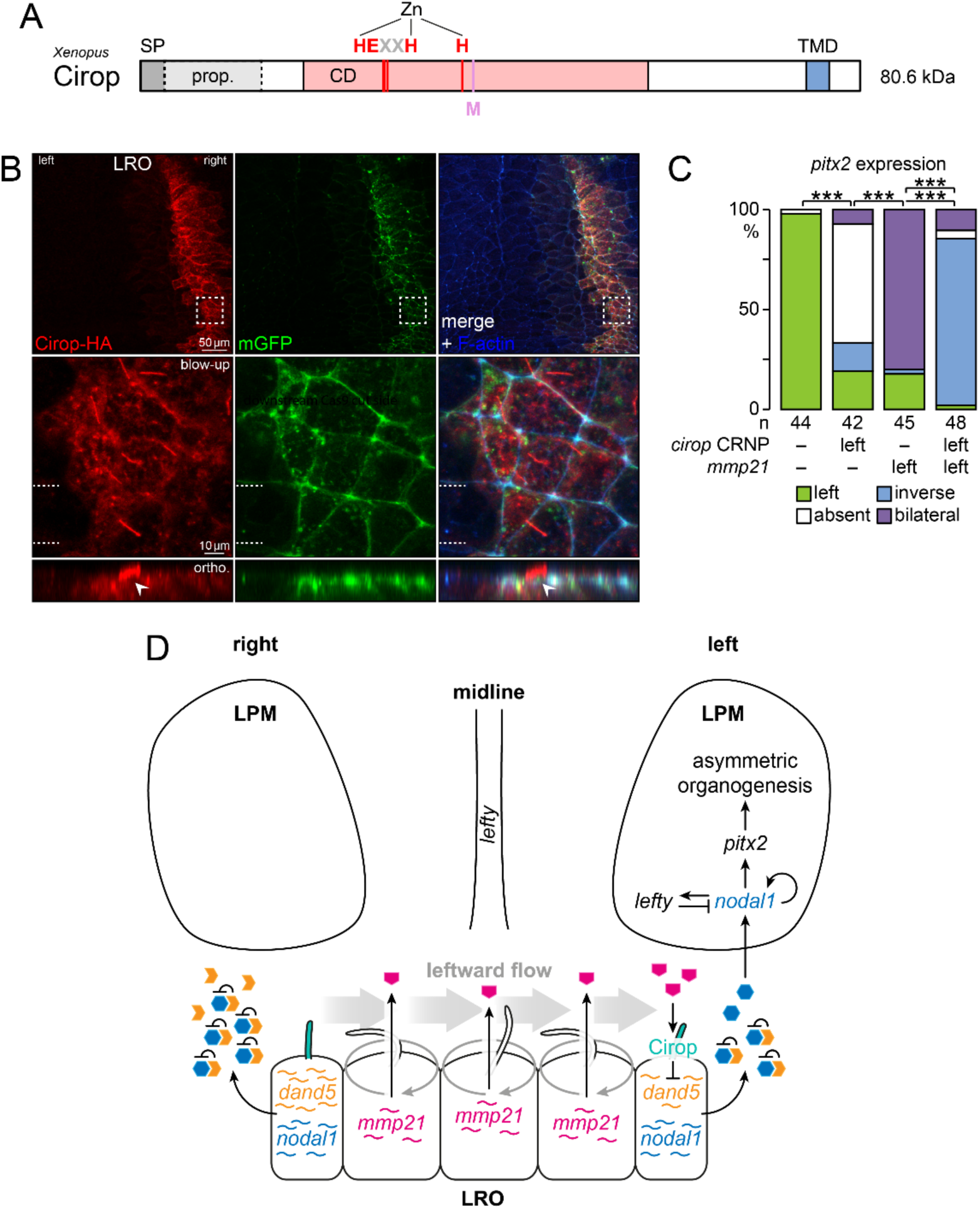
Cirop is a downstream transducer of Mmp21 at LRO cilia. **(A)** Cirop is composed of an N-terminal signal peptide (SP), a propeptide, a catalytic domain (CD) and a C-terminal transmembrane domain (TMD). Coordination of the Zn atom as well as the conserved residues of the catalytic site (HEXXH plus C-terminal H) and methionine-turn (M) are depicted. **(B)** C-terminally HA tagged Cirop localizes to LRO cilia and accumulates at cell borders. Cirop-HA producing cells are marked by co-expressed mGFP. Cell borders are visualized by F-actin staining. Dashed white line squares outline the magnified regions, highlighting the ciliary localization of Cirop-HA. An orthogonal view of the region, indicated by two dashed white lines, shows a single protruding cilium marked by a continuous apical Cirop-HA signal (white arrowheads). **(C)** Cirop acts downstream of Mmp21, as loss of the Nodal cascade in left-sided *cirop* f0 mutants can be transformed into right-sided cascade activity by co-injecting the mRNA of diffusible Mmp21. **(D)** Model showing the key steps of laterality determination in vertebrates with a fluid flow generating LRO: Central LRO cells generate a cilia-driven leftward fluid flow and secrete Mmp21 from their apical side. Flow directs diffusion of Mmp21 to left-sided LRO sensor cells. Sensor cells perceive Mmp21 via Cirop-positive sensory cilia. Cirop-dependent signaling of Mmp21 represses *dand5* on mRNA level, attenuating Dand5 protein levels on the left side. Left-sided Nodal1 is released from repression by Dand5 and induces the Nodal signaling cascade in the left lateral plate mesoderm (LPM).

To test the hierarchical placement of these two peptidases, *mmp21* mRNA and *cirop* CRNPs were co-injected into left blastomeres, leading to left-sided *cirop* mutants that simultaneously overexpress Mmp21 on the ipsilateral side (Fig. 4C). In these embryos, Mmp21 was unable to rescue the absence of the Nodal cascade caused by left-sided loss of Cirop. Instead, Nodal signaling was largely inverted, which is in line with the diffusible nature of Mmp21 and its placement upstream of Cirop, which remained unedited and thus functional on the right side.

In summary, our analysis uncovered the underlying pathomechanism of HTX, attributed to homozygous *MMP21* null variants like the novel p.(L180Gfs*74) variant described in this study. It revealed that Mmp21 conveys the positional information for the hitherto unspecified left side as a fluid flow transported morphogen.

## Discussion

The number of reported *MMP21* variants associated with HTX is increasing, but the function and hierarchical position of Mmp21 within the underlying process of laterality determination remained enigmatic. Our functional study shows that Mmp21 has a pivotal role in symmetry breakage and builds a functional unit with the leftward fluid flow. We would like to propose that Mmp21 embodies the long sought-after instructive cue, enriched asymmetrically by the ciliated LRO.

Our model suggests the following steps (Fig. 4D): (1) Central LRO cells synthesize and release Mmp21 from their apical side. (2) The cilia driven leftward fluid flow guides the diffusion of Mmp21 towards left LRO sensor cells. (3) Mmp21 signaling through ciliary localized Cirop in left sensor cells eventually triggers the post-transcriptional downregulation of *dand5*, leading to the activation of Nodal signaling on the left side.

This model is a derivation of the so-called morphogen model, originally suggested in the groundbreaking paper by the Hirokawa group on the identification of leftward fluid flow^5^. It provides a compelling explanation for the *mmp21* LOF and GOF results across many levels of analysis. The Nodal cascade pattern provides a useful readout to assess the rigor of mechanistic models in the laterality field^32^, despite not being analyzed or discussed in many studies nowadays. In the case of Mmp21 LOF, absence of the Nodal cascade can be attributed to the loss of the instructive signal, namely Mmp21, carrying the identity information for the left side. In contrast, the bilateral Nodal cascade observed after Mmp21 GOF results from excessive signal, which has the ability to diffuse against the fluid flow under non-physiological conditions. The Hirokawa group postulated that only proteins in the range of 20-40 kDa could generate a stationary gradient over the LRO^6^. We interpolated their data and concluded that it is valid to extend this to 50 kDa, matching the molecular weight of mature Mmp21 (Fig. S1A). However, as Mmp21 resembled the localization of Wnt and Hedgehog ligands during their proteoglycan-associated spread in naïve epithelia cells^29,30^, one can speculate that interactions with components of the extracellular matrix may occur. Such interactions could limit arbitrary diffusion of Mmp21 within the flow medium in accordance with absence of the Nodal cascade in cases of disturbed cilia motility^33^.

Our model contrasts with the so-called two-cilia model, which is based on Brueckner and colleagues’ idea that mechanosensory cilia directly detect the fluid flow *per se*^7,34^. The Hamada and Yuan groups independently demonstrated that bending of LRO cilia induces *dand5* flow response in sensor cells, using an elaborate optical tweezer setup^35,36^. This response was linked with a higher number of intracellular calcium ion transients and depended on Polycistin 2 (Pkd2), which forms homo-^37^ or heterotetrametric cation channels^38^.

Interestingly, Pkd2 complexes with the potential channel subunit Polycistin 1 like 1 (Pkd1l1) presumably on LRO cilia^39^. Like *MMP21* and *CIROP*, *PKD1L1* is associated with HTX^40^ (MIM 617205) and belongs to the same genetic module that is only preserved in species with a flow generating LRO^3^. As Pkd1l1 appears to be an upstream repressor of Pkd2^41^, one can speculate that Mmp21 and Cirop directly inactivate Pkd1l1 through its large extracellular domain, linking protease activity to flow-related calcium signaling.

Of note, we do not exclude the possibility that bending of LRO cilia has instructive capacity for the left-right axis. A scenario is conceivable in which the basic mechanisms from the two-cilia and morphogen model cooperate to increase systemic robustness, thus lowering the intrinsic frequency of laterality defects. Such a scenario is also supported by a recent report from the Niehrs group published during the preparation of this manuscript^49^. The authors describe that the secreted growth factor R-spondin 2 (Rspo2), which has not yet been described to be causative for human laterality defects, appears to act as a fluid flow transported morphogen in *Xenopus* as well. Deciphering the hierarchy of these potentially interlinked determinants in different model organisms will shed light on the composition of the core module required for leftward fluid flow perception.

## Methods

### Exome sequencing

Exome sequencing was employed for the detection of variants in the female proband II-6 (ID 4263)^3^. One microgram of high-quality gDNA was used for exome capture with the Ion TargetSeq Exome Kit. The exome library was prepared on an Ion OneTouch System and sequenced on an Ion Proton instrument (Thermo Fisher Scientific) using one Ion PI chip. Sequence reads were aligned to the human reference genome (GRCh37/hg19) using Torrent Mapping Alignment Program from the Torrent Suite v5.0.2. The variants were called using the Torrent Variant Caller plugin v5.0.2 and were annotated with the associated gene, location, protein position and amino acid changes, quality-score, coverage, predicted functional consequences using protein position and amino acid changes, Sorting Tolerant From Intolerant (SIFT)^42^, Polymorphism Phenotyping v2 (PolyPhen-2)^43^ and Grantham^44^ prediction scores, phylogenetic p-values (PhyloP)^45^ conservation scores and 5000 genomes Minor Allele Frequencies. Variants were filtered for common single nucleotide polymorphisms using the NCBI’s “common and no known medical impacts” database (ftp://ftp.ncbi.nlm.nih.gov/pub/clinvar/vcf_GRCh37/), the Exome Aggregate Consortium (ftp://ftp.broadinstitute.org/pub/ExAC_release/release0.2/) and the Exome Sequencing Project (http://evs.gs.washington.edu/EVS/). We next removed variants that were present in greater than 1 % of the previously in-house 465 sequenced samples. A total of 15.8 Gb was sequenced with an average read length of 176 bp. An average coverage of 186x was achieved over the exome, with 97 % of bases covered at least 20x for each individual. A combined total of 52 438 variants were identified across protein-coding exons, untranslated regions, splice sites and flanking introns. Additional filters were applied to retain exonic and splice variants that were homozygous in the proband. A final set of 17 variants remained, of which 8 were in conserved amino acids (PhyloP score > 1). Within those 8 variants, the biallelic variant c.537_547delGCTGCTGGGCG, p.(L180Gfs*74) in gene *MMP21* was identified.

### Protein structure prediction

The Mmp21 (NM_001085816.1) amino acid sequence C-terminal of the FCS was used as sequence query in a ColabFold^23^ run with standard settings. The highest rank model was considered highly confident with a predicted local distance difference test (pLDDT) score of 90.9 and a predicted template modeling (pTM) score of 0.885. AlphaFill^46^ was subsequently employed to enrich the model with the cofactor zinc.

### Experimental animals

The treatment of *Xenopus laevis* frogs (Nasco) was in accordance with the German Animal Welfare Act and was approved by the Regional Government of Stuttgart, Germany. *Xenopus* females were primed for spawning by injecting 50 units human chorionic gonadotropin (CG, Merck) 3-6 days before embryos were needed. Collection of oocytes began 12 h after injecting additional 300-600 units GC to induce ovulation. *Xenopus* males were sacrificed and testis were isolated to obtain sperms for subsequent *in vitro* fertilization.

### Intracytoplasmic injection experiments

Microinjections were performed at the 1- or 4-cell stage using the following reagents and doses in volumes of 4 or 8 nl: 1 ng *Streptococcus pyogenes* Cas9 with NLS (PNA Bio) plus 200 pg *mmp21* sgRNA (5’-GGTGATGGTGGCATCATCAA-backbone-3’) or 200 pg *cirop* sgRNA (5’-GGGGAAGTCAGGGATCTAGG-backbone-3’), 0.75 pmol *dand5* TBMO (Gene Tools, 5’-TGGTGGCCTGGAACAGCAGCATGTC-3’), 100-400 pg C-terminal HA-tagged or untagged wild-type Mmp21 mRNA, 120 pg single or triple mutant Mmp21 mRNA, 200 pg C-terminal HA-tagged Cirop mRNA and 200 pg mGFP mRNA.

CRISPRscan^47^ was utilized for sgRNA design and the corresponding DNA templates were created in an oligo extension reaction with Pfu DNA polymerase (Promega). Templates were cleaned with the GeneJET Gel Extraction and DNA Cleanup Micro Kit (Thermo Fisher Scientific). The T7 MEGAshortscript Kit (Thermo Fisher Scientific) was combined with the MEGAclear Transcription Clean-Up Kit (Thermo Fisher Scientific) for sgRNA synthesis and purification. Genome editing of *mmp21* was analyzed via direct sequencing of PCR products^24^ generated with target site flanking forward (5’AGTCTGATGCAAGAGGGAGT3’) and reverse (5’TTGCCTGTGGATTTCTGAAGC3’) primers on pooled genomic DNA from stage 45 specimens. Mutagenesis rates were determined with Synthego ICE (https://ice.synthego.com/).

All mRNAs were transcribed with the SP6 mMESSAGE mMACHINE Kit (Thermo Fisher Scientific) and cleaned-up with the GeneJET RNA Purification Kit (Thermo Fisher Scientific).

### Archenteron injections

MC was diluted in 1x Modified Barth’s Solution (MBS) to 2 % and paraxially injected into the archenteron between stage 14 and 15. Control embryos were injected with 1x MBS without MC. After the injection, the specimens were incubated at 26 °C until reaching stage 19, in order to enhance the viscosity of the MC solution^31^.

### Brefeldin A incubation

Brefeldin A (Selleck Chemicals) was diluted to 1 mM stocks in dimethyl sulfoxide (DMSO). Working solutions of 2 nM were generated by a serial dilution with 0.1x MBS. *Xenopus* embryos were incubated from stage 8 until stage 17 and subsequently processed for fluorescence staining.

### RNA *in situ* hybridization (ISH) and fluorescence staining

Embryos were fixed overnight in 100 mM MOPS, 2 mM EGTA, 1 mM MgSO4 and 3.7 % formaldehyde (MEMFA) and processed according to standard procedures. Antisense RNA ISH probes were synthesized with DIG RNA Labeling Mix (Roche) using T7 or SP6 RNA polymerases (Promega). Probes were detected with alkaline phosphatase (AP) conjugated anti-DIG Fab fragments (Roche). BM-Purple (Roche) was used as AP substrate. Localization of HA tagged proteins were analyzed with an anti-HA tag antibody (Roche, clone 3F10, 1 : 500) and Alexa Fluor coupled secondary antibodies (Thermo Fisher Scientific, 1 : 500). F-actin was visualized via Alexa Fluor 405 phalloidin (Thermo Fisher Scientific, 1 : 100).

### LRO Analysis

Leftward fluid flow was monitored on dorsal explants of stage 17 embryos in a 30 s time window using 0.5 µm yellow-green FluoSpheres (Thermo Fisher Scientific, 1 : 2500). Flow speed and directionality were quantified with the ImageJ plugin Particle Tracker^48^ and a custom-made script written in R^31^. Sizes of the *nodal1* and *dand5* expression domains were determined with the Color Threshold tool of ImageJ^48^.

### Statistics

Probability values were calculated for metric data with the two-tailed Wilcoxon signed-rank test and for ordinal data with the two-tailed Fisher’s exact test. Significance levels were defined as p ≥ 0.05 is not significant, p < 0.05 is 1 asterisk, p < 0.01 is 2 asterisks and p < 0.001 is 3 asterisks. The number of analyzed embryos is given by n.

## Acknowledgments

We thank Jennifer Kreis for the critical proofreading of the manuscript. Furthermore, we would like to thank Sumanty Tohari, Alvin Yu-Jin Ng and Byrappa Venkatesh for assistance with exome sequencing.

## Author contributions

T. O. designed the study, supervised experiments and wrote the manuscript.

T. O., A. B., Y. K. and O. A. performed experiments.

E. S.-R. and B. R. performed and supervised exome sequencing and helped in manuscript editing.

M. T. validated ordinal data blindly.

M. B. and A. S. provided laboratory resources and infrastructure.

S. W. and P. B. made clinical diagnoses and collected clinical data and samples.

---

**Fig. S1:**
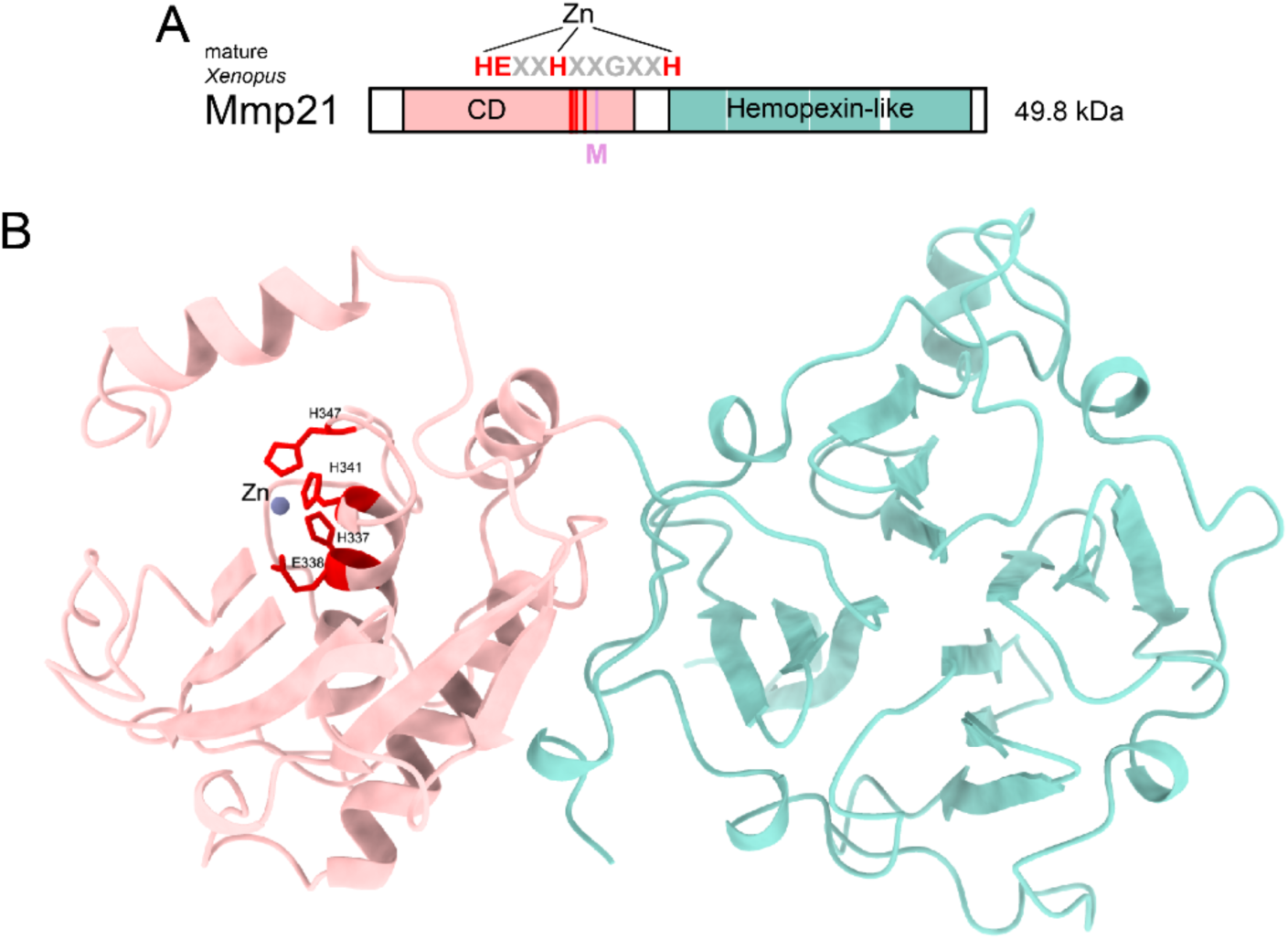
Mature Mmp21 has a predicted tertiary structure with two distinct globular domains. **(A)** Removal of the propeptide lowers the molecular weight of Mmp21 to 49.8 kDa. **(B)** ColabFold was employed to predict the conformation of mature Mmp21, which adopts a bipolar, globular structure formed by the catalytic domain (CD) and the Hemopexin-like beta-propeller. AlphaFill enriched the model with the catalytic zinc atom. The conserved histidine and glutamate residues (H337, E338, H341 and H347) of the catalytic core are displayed with their side chains in red. All other residues N-terminal of the C-terminal CD border were colored in light red, while residues C-terminal of the CD were colored in turquoise. Visualization of the catalytic cleft and core was optimized by concealing the loop between A304 and D311.

**Fig. S2:**
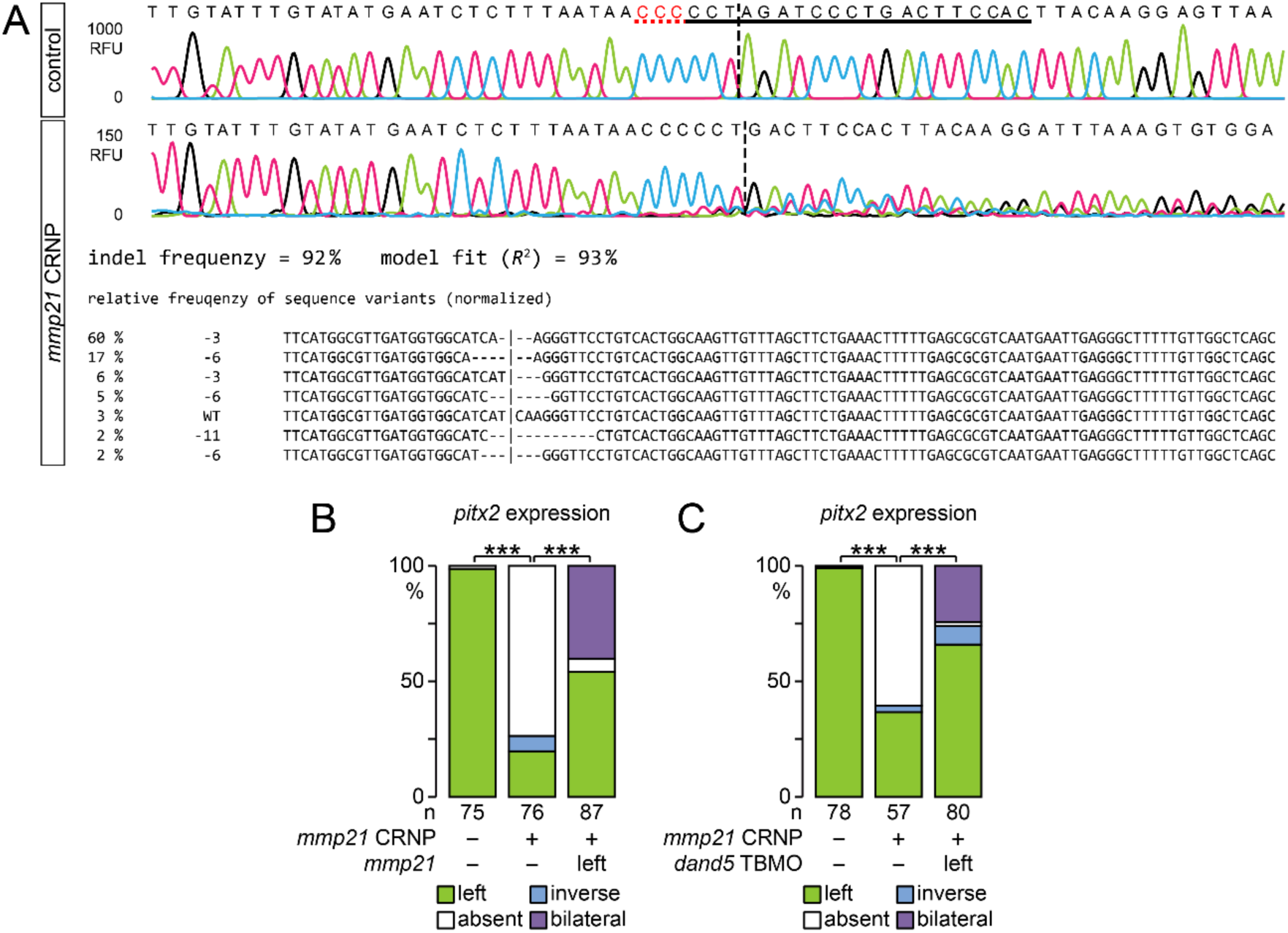
Mutagenesis of *mmp21* is efficient and creates rescuable laterality phenotype. **(A)** Sanger sequencing data of the wild-type and mutagenized *mmp21* region is displayed as chromatogram. Signal intensity is given in relative fluorescence units (RFU). Synthego ICE analysis confirms efficient mutagenesis, resulting in various genomic DNA species with small deletions. The sgRNA target site is underscored in black and the protospacer adjacent motif with a dashed red line. Vertical black dashed lines indicate the site of the Cas9-induced DNA double-strand break. **(B)** Specificity of the *mmp21* crispant phenotype was verified by the rescue potential of *mmp21* mRNA, which restored left-sided *pitx2* expression. **(C)** Expression of *pitx2* was also rescued by a left-sided knockdown of *dand5*, mediated by a translation blocking morpholino oligomer (TBMO), showing that events downstream of flow-dependent *dand5* repression are Mmp21 independent.

**Fig. S3:**
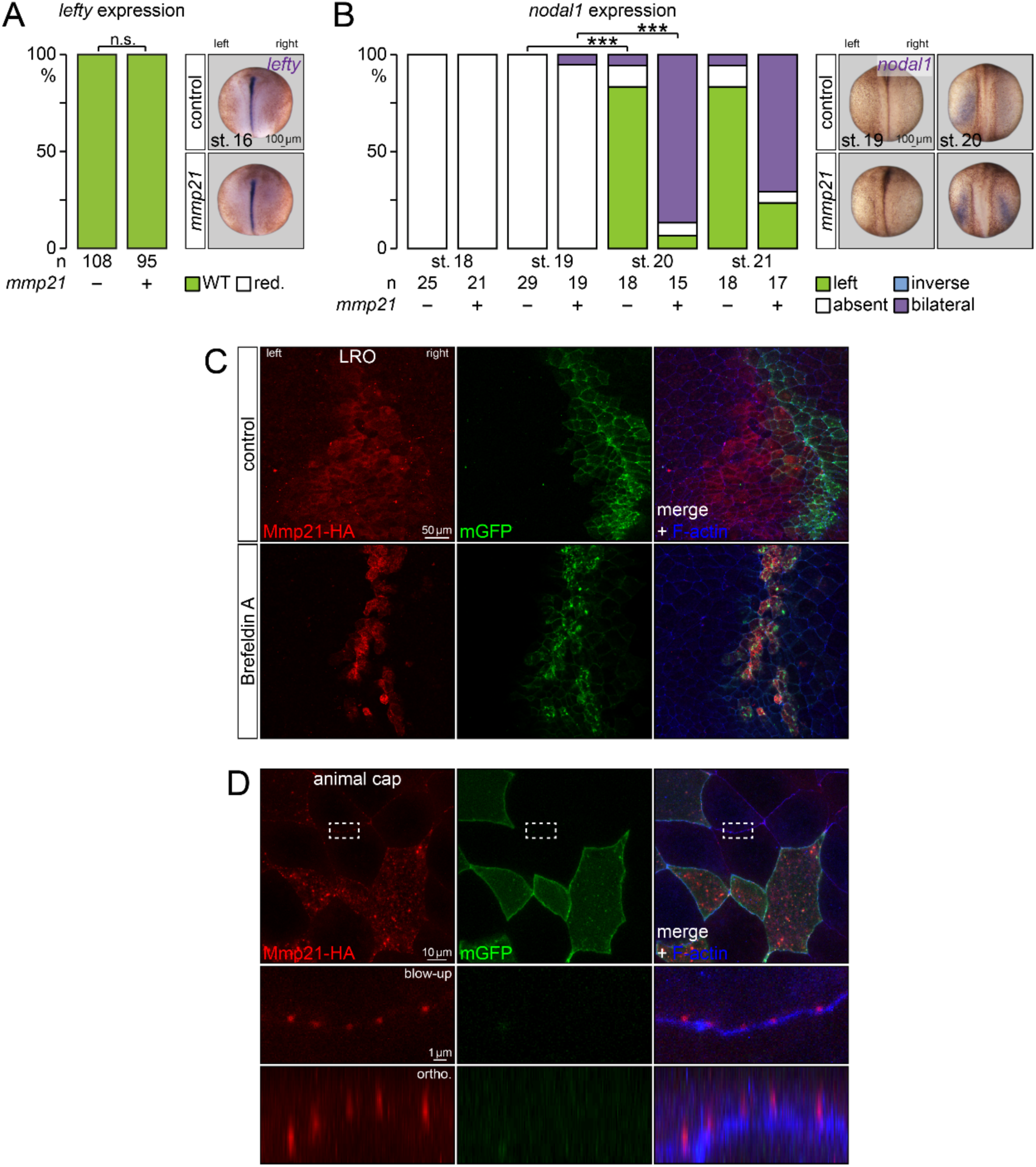
Secreted Mmp21 does not affect the midline barrier. **(A)** Overexpression of Mmp21 does not reduce the mRNA level of *lefty* in the midline. **(B)** Bilateral Nodal cascade activity, caused by elevated Mmp21 levels, is not the result of midline leakage, as *nodal1* expression started simultaneously between the left and right side. **(C)** Inhibition of secretion via Brefeldin A incubation prevents spreading of Mmp21-HA over LRO tissue. Mmp21-HA synthesizing cells are indicated by mGFP, which was used as lineage tracer. Cell borders are marked by F-actin. **(D)** Diffusion of Mmp21 in LRO-unrelated naïve epithelial cells occurs along cell borders, which are visualized by F-actin. Cell bodies and borders of Mmp21 secreting cells are marked by co-expressed mGFP. Dashed white line rectangles outline the magnified regions, highlighting Mmp21-HA foci between non-secreting cells. Orthogonal view shows apical aggregation of Mmp21-HA.

